# Genetic parameters and sex-specific architecture of observed and latent fertility phenotypes in a closed breeding nucleus of an Arctic salmonid

**DOI:** 10.1101/2025.10.16.682826

**Authors:** Fotis Pappas, Martin Johnsson, Paul Vincent Debes, Christos Palaiokostas

## Abstract

Successful reproduction is a key factor for efficient breeding schemes and sustainable animal farming. Aquaculture breeding programs rely heavily on small fractions of selected breeders to yield large production stocks, given the high fecundity typically observed in these species. In Sweden, Arctic charr (*Salvelinus alpinus*) is a salmonid with notable commercial potential, with a selective breeding program operating for ten generations under a growth-rate focused breeding goal. Despite significant gains, the nucleus faces challenges with low and fluctuating fertility impeding expansion efforts. In this study, we estimate genetic parameters for charr milt quality phenotypes measured with specialized cytometry and Computer-Assisted Sperm Analysis (CASA). At the same time, we assess the sex-specific architecture governing egg count and sperm concentration along body size. Finally, we propose a novel analytical framework for the analysis of realized fertilization success rates by considering a multiplicative system of latent maternal and paternal contributions. Low to moderate heritability estimates and genetic correlations were obtained from multi-trait modelling for traits reflecting sperm quality along with high estimates for fork length. Genetic correlations among sperm kinematic parameters appeared strong, while the same traits showed weak positive and weak negative correlations with sperm concentration and fork length, respectively. Furthermore, a negative genetic correlation between sperm concentration and both male body size and egg count suggests a complex interplay of a possible trade-off and sexual antagonism. Our latent fertility analytical approach returned low to moderate heritability estimates depending on the modelling configuration. Overall, our study demonstrated the complexity characterizing the heritable portions of reproductive traits in Arctic charr and tested alternative tools that have the potential for integration into selective breeding programs.

## Introduction

Fertility is a complex phenotype governed by multiple genetic and environmental factors. Sexual reproduction entails dependence on the biological properties of more than one individual, making reproductive success a function of phenotypic combinations. Rather than being a single biological trait, the ability to reproduce reflects the interaction of various female and male elements such as hormonal regulation (Findlay et al., 1992; Santhakumar, 2024), reproductive tract health and gamete quality (Bobe, 2015; Cao et al., 2024; Clarkson et al., 2024; Fernández-López et al., 2022; Kumaresan et al., 2020). The latter includes, but is not limited to egg size, sperm motility, energy status, morphology, and genetic material integrity. In wild populations, fertility is considered a direct indicator of evolutionary fitness, with reduced reproductive rates often pointing to inbreeding depression or challenging environmental conditions. Inbreeding, by increasing homozygosity, can expose unfavourable recessive alleles that negatively impact reproductive traits (Charlesworth & Willis, 2009), from gametogenesis to embryonic viability and contribute to demographic decay.

Domesticated animal populations undergo strong artificial bottlenecks for traits of economic importance, such as growth-rate, through selective breeding. However, genetic trade-off patterns between production and reproduction have been detected in multiple livestock species, often revealing an unfavourable relationship between mobilization of resources for biomass and reproductive investment (Akanno et al., 2013; Andersen-Ranberg et al., 2005; Piles & Tusell, 2012). On top of that, reproduction in captivity is most often carried out through controlled and restricted artificial mating events (can be assortative), sometimes with no actual contact of the breeders (e.g. with artificial insemination), thereby introducing additional sources of variation due to husbandry and handling.

In aquaculture species, the success and profitability of breeding programs depend partly on using only a small number of selected breeders to generate production stocks that are orders of magnitude larger (Gjedrem et al., 2012). High fecundity is therefore a prerequisite to make such breeding schemes viable. This high reproductive output, combined with external fertilization and a controlled, homogenous environment makes farmed fish an excellent model for studying fertility. Aquaculture settings enable detailed assessment of fertility factors in both male and female breeders, thanks to macroscopic observations from artificial mating to post-hatching developmental stages. In salmonids specifically, fertilization is achieved only through human input and hatching of the eggs occurs in a biosecure environment that allows the disentanglement of risk factors and, when applicable, the identification of genetic influences on reproductive success. Despite the existence of strong evidence for unfavourable genetic correlations between early maturing males (occurring even when the animals are less than one year old) and growth rate in salmonid species (Gjerde, 1984; Martyniuk et al., 2003; Tsai et al., 2015), the respective genetic and phenotypic relationships of production traits with on-farm reproductive success, such as between growth rate and fecundity or milt quality, remain understudied.

Arctic charr (*Salvelinus alpinus*) is a salmonid fish with desirable gastronomical characteristics and consequently high economic value. The species is cold-adapted and has been successfully introduced to aquaculture in countries neighbouring the Arctic (Sæther et al., 2013). In Sweden, the oldest selective breeding program operates for approximately 40 years (10 non-overlapping generations) using Arctic charr of landlocked origin and has achieved notable genetic gains for growth rate (Carlberg et al., 2018). Though reduced and varying reproductive performance of that broodstock introduces scaling challenges (Palaiokostas et al., 2021). Preliminary investigations suggest a major paternal factor in realized reproductive success (Jeuthe et al., 2019) that could be partly explained by heritable factors as suggested by a study on milt quality (Kurta et al., 2022).

In this study, we utilize reproductive success, fecundity and milt quality data from the Swedish Arctic charr (Arctic Superior®) breeding program to estimate genetic parameters of male and female fertility proxies. Furthermore, we attempted to shed light on sex-specific architecture and genetic correlations between fecundity and growth. Finally, we propose a novel modelling approach that considers the probability of fertilization success to be a product of latent fertility coefficients for sires and dams.

## Materials and methods

### Retrospective fecundity and fertilization rate data

A retrospective dataset was utilized, consisting of 612 observations from artificial mating events involving 395 sires and 544 dams. The artificial matings were carried out for the resetting of the non-overlapping generations in the Swedish national charr breeding program. For each mating with known dam and sire identities, counts of spawned and eyed eggs were recorded. All breeders had corresponding pedigree records, with the exception of 53 dams that were treated as unrelated and non-inbred phantom animals. Additional features such as year, fork length and body weight were also available. In total ∼3.9 million spawned eggs were recorded, of which ∼1.1 million were successfully fertilized.

### Sperm quality parameters

A total of 1,106 male charrs from the 8^th^ (N=370) and 9^th^ (N=736) generations of the breeding program were milted and their fork length was measured (mm, ±10 mm) during the 2020 and 2024 spawning seasons, respectively. Fish were reared in indoor concrete tanks with a flow-through water supply and ambient water temperature. After anaesthetizing the fish using MS-222 (Sigma-Aldrich, Darmstadt, Germany), milt samples were collected in sterile containers and stored at 4 °C. A set of sperm quality phenotypes was measured within six hours post-sampling. More specifically, sperm motility was assessed using a computer-assisted sperm analysis (CASA) setup coupled with the SCA® Motility imaging software (Microptic S.L., Barcelona, Spain) as previously described (Kurta et al., 2022). CASA measurements for each sample were taken two to three times using 20 µm depth slides with two counting chambers (CellVision, Heerhugowaard, The Netherlands). Milt activation was achieved using either water (2020 trial) or the fish sperm activation solution ActiFish® from IMV technologies (2024 trial).

At 15 s post-activation, the following metrics were extracted: curvilinear velocity (VCL μm/s), proportion of rapid spermatozoa (PRS) and straightness of trajectory (STR, the ratio between straight line velocity and average path velocity), while late curvilinear velocity (LVCL μm/s) was recorded at 30 s. Records for the latter two traits were available only for the males of the 9^th^ generation. A frame rate of 100 fps (50 frames) was used together with classification thresholds of VCL ≥ 20 µm/s for motile and VCL ≥ 100 µm/s for rapidly motile spermatozoa. Finally, sperm concentration (SC, ×10^6^ cells/mL) was measured with sperm-specific cytometry using NucleoCounter® SP-100™ (ChemoMetec A/S, Allerod, Denmark).

### Genetic parameters for sperm quality parameters and growth

A multi-trait approach was employed in the case of sperm quality parameters and growth for male charrs using the BLUPF90 suite (Lourenco et al., 2023; Misztal et al., 2022). Six traits were analysed, namely sperm concentration (SC), curvilinear velocity recorded at 15 s post-activation (VCL), late curvilinear velocity measured at 30 s (LVCL), proportion of rapid spermatozoa (PRS), straightness of trajectory (STR) and male fork length (FL(m)). These traits were modelled using animal models that accounted for the fixed effect of trial year and the covariate of water temperature on the day of sampling (the latter applied only to sperm traits).

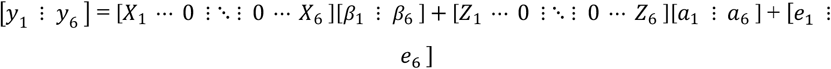

Phenotype vectors ***y*** are expressed as functions of the aforementioned fixed effects (vectors *β*) via the design matrices ***X***, additive genetic effects 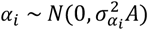 through design matrices ***Z*** and residuals 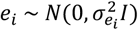 . Matrices ***A*** and ***I*** denote the additive genetic relationship and identity matrix, respectively.

Additionally, genetic marker information (7,939 SNPs) was available for 1,049 animals from the closed nucleus (Palaiokostas et al., 2022). A subset of 271 animals, corresponding to males of generation 8 with sperm records (∼1/4^th^ of all phenotyped individuals), provided a link for using the single-step methodology (ssBLUP) under the same model structure (Multi-trait Model 1). The input data were standardized to reflect z-scores and 150 initial rounds of the expectation maximization algorithm (EM-REML) were used for variance components estimation (i.e. the 6×6 variance-covariance matrices).

### Genetic parameters for female fecundity and covariance with sperm concentration and growth

In a similar fashion to the above, a multivariate model (Multi-trait model 2) was fitted for the female-restricted phenotypes egg count (EC) and female fork length (FL(f)) from the retrospective dataset in combination with male-restricted traits SC and FL(m). The aim was to obtain insights into the genetic architecture governing fecundity and male reproductive potential in relation to male and female growth rate. As above, the trial year was accounted as a fixed effect for all four traits, while female fork-length (FL(f)) was treated as a covariate of egg count (EC) and water temperature on the day of sampling as a covariate for sperm concentration (SC).

### Bayesian hierarchical framework for analysis of latent fertility coefficients

The proportion of fertilized fish eggs (reaching the eyed egg stage) can be expressed as a function of male and female fertility. The goal is to separate male and female reproductive contributions to the outcome of each cross by incorporating the known structure due to additive genetic relationships and environmental effects. We therefore model the number of eyed eggs from mating *i* (sire *m*_*i*_, *dam f*_*i*_) as:

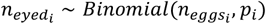

where *p*_*i*_ is the probability that an egg in cross *i* gets fertilized and reaches the eyed stage.

#### Fertilization model 1

In the simplest form, fertilization success is the product of a female fertility component and a male fertility component:

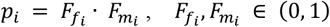

*F*_*f*_ is the latent female fertility coefficient reflecting the proportion of viable, fertilizable eggs and *F*_*m*_ is the latent male fertility corresponding to the ability of the ejaculate to fertilize viable eggs (*F*_*f*_ · *n*_*eggs*_) with *n*_*eggs*_ being the total number of eggs spawned by the dam. Female and male components are latent liabilities mapped to (0, 1) via a probit link:

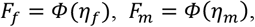

where Φ is the standard normal cumulative density function (CDF).

In matrix notation and given the additive genetic relationship matrix ***A*** built with the R package AGHmatrix v2.1.4 (Amadeu et al., 2023), the linear predictors *η*_*f*_ and *η*_*m*_ were modelled as such:

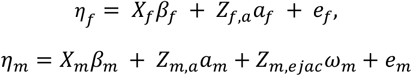

Where *X* the design matrices for fixed effects *β* (intercept and year for females; intercept, year and use of sperm boosting solution for males), *Z*_*f,a*_ and *Z*_*m,a*_ the incidence matrices for female and male additive genetic effects 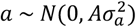 modelled in a bivariate fashion and 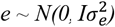 are the residual terms. In the case of males, an ejaculate permanent environment effect 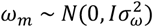 was added to account for heavy sire replication (same ejaculate used for more than one dam). In all cases, *I* denotes the identity matrix.

#### Fertilization model 2

The above model assumes that male fertility is independent of the number of eggs spawned by the female (Figure 1A). However, this assumption might not be biologically realistic in many cases and especially considering highly fecund and externally fertilizing systems. In the latter, ratios of potent spermatozoa to eggs most probably influence fertilization outcomes. In reality, a male’s capacity to fertilize may be mainly constrained by sperm concentration, motility and milt volume, especially when paired with highly fecund females.

**Figure 1:**
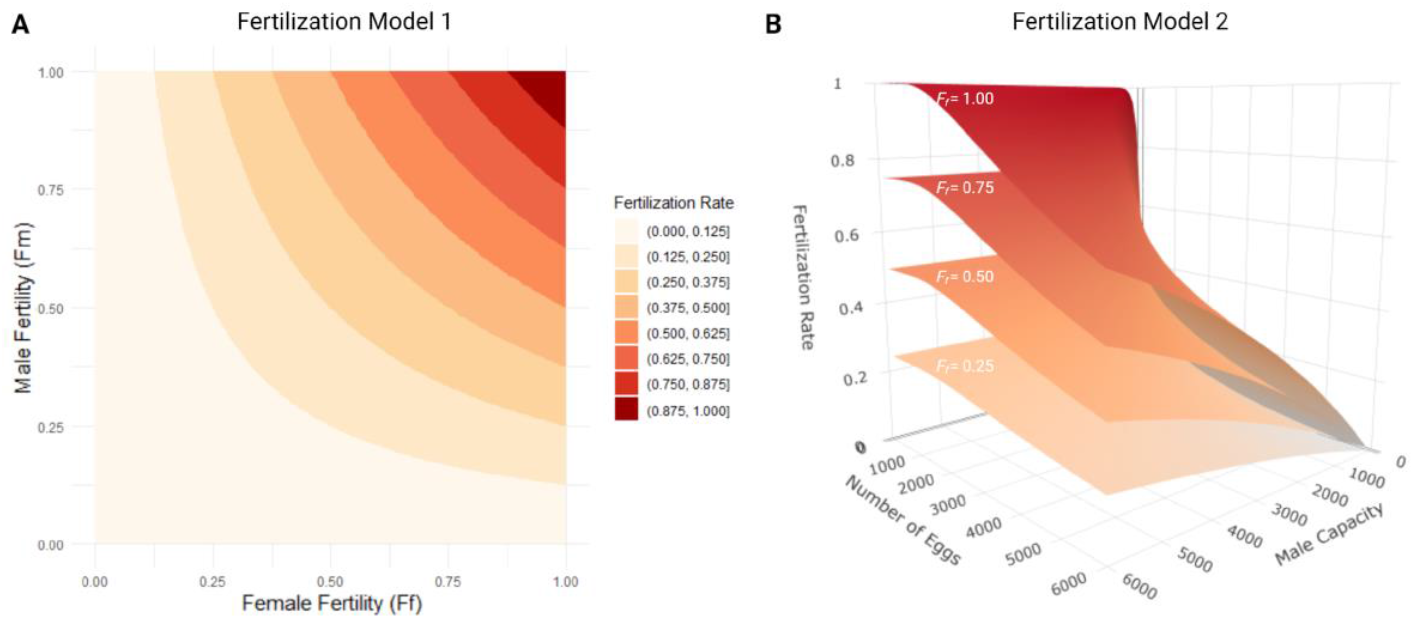
Color-coded contour visualizations of fertilization rate (FR) as a function of underlying fertility parameters. **A)** Contour plot for Fertilization Model 1, where FR is considered a product of latent male and female fertility coefficients *F*_*m*_ and *F*_*f*_. **B)** Surface plot for Fertilization Model 2, where FR is expressed as a non-linear function of *F*_*f*_, male fertilization capacity *C*_*m*_ and number of spawned eggs. Surfaces for *F*_*f*_ values fixed at 0.25, 0.50, 0.75 and 1.00 are presented. Figure was created with ggplot2 v3.4.4 and BioRender.com.

To address this, we consider a second model formulation where male fertility is represented by a fertilization capacity parameter, *C*_*m*_ > 0 and fertilization success depends on the ratio of male capacity to female fecundity (reflecting available fertile spermatozoa per egg). This saturating model allows for the biologically realistic scenario where males may fully fertilize small spawns, but only partially fertilize larger ones, thus incorporating the asymptotic relationship between sperm availability and number of spawned eggs. In this case, we define:

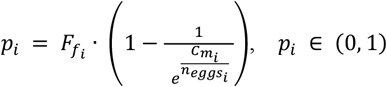

In practice, this second fertilization model assumes that sperm-egg interactions can be approximated by a Poisson process, where the probability of fertilization increases with the abundance of potent spermatozoa per egg (Figure 1B). The factor 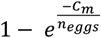 corresponds to the probability of at least one successful sperm collision per egg under a Poisson process. This is a simplified version of similar biophysical functions that have been previously proposed for fertilization saturation in free spawning species (Styan, 1998; Vogel et al., 1982). Sub-models for *F*_*f*_ and *C*_*m*_ were formulated in the same way as in Model 1 except that the male term *C*_*m*_ now is mapped with a log link (not a probit link), whereas the female liability remains on the probit scale.

In this model, in addition to fertilization rate, number of eggs is also necessary. Due to lack of such records for a subset of 62 mating events that had a fertilization rate of 0, imputed egg numbers were used. Imputation took place using the mice v3.17.0 (Buuren & Groothuis-Oudshoorn, 2011) R package using dam weight, dam length and year as features.

Weakly-informative, realistic priors were assigned to parameters of both models (https://github.com/pappasfotios/AcFertilityGP/tree/main/FertRate_Models). Specifically, intercepts for male and female fertility were centered at 0.5 (probit scale), while sperm capacity (*C*_*m*_, Fertilization model 2) was assigned a tighter log-rate intercept prior implying near-saturating capacity for typical clutch sizes (mean=6,439, sd=2,717.44). Genetic and other random effect priors encouraged data-driven, moderate heritability estimates (inspired by variance components of fertility proxies) while discouraging strong genetic dependence among traits. Implementation was carried out using Stan via R/rstan (R Core Team, 2023; Stan Development Team, 2023) and 4 chains of 10,000 iterations, out of which 4,000 were used for warmup. Model convergence was assessed with visual inspection of trace plots and by assessing metrics like 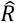 (indicator of convergence of the Markov chain to the target distribution) and the effective sample size (ESS).

In both models, heritability estimates for the latent variables of female fertility are given by:

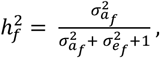

while for male fertility we have:

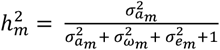 or 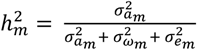 depending on whether that is expressed on the probit or log scale (de Villemereuil et al., 2016) as in Fertilization Model 1 and Fertilization Model 2, respectively.

## Results

### Estimation of variance components for sperm quality parameters and covariance with growth

Multi-trait analyses for male phenotypes yielded low to moderate heritability estimates for all milt traits and a higher estimate for fork length (Table 1). More specifically, heritability ranged from ∼ 0.1 to ∼ 0.3 for sperm parameters and ∼ 0.6 for fork length. Those estimates were consistent across the two approaches (pedigree BLUP and single-step BLUP), with the only actual deviations observed for SC and FL, although these were within the range of overlaps defined by the respective standard errors.

**Table 1.**
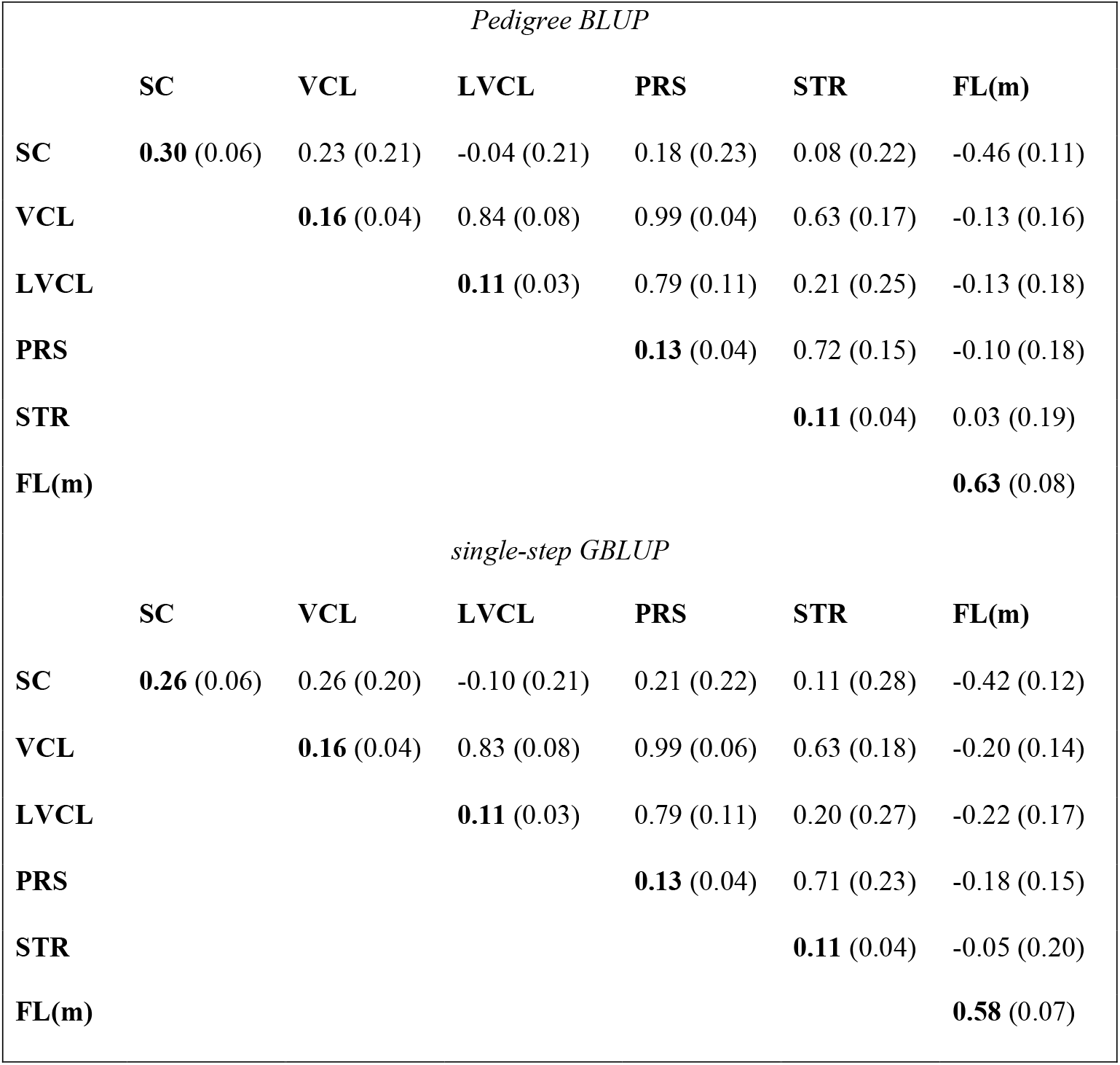
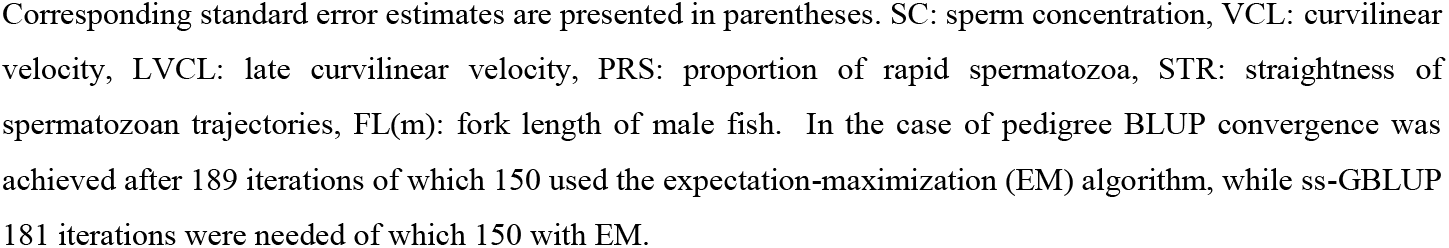
Heritability (diagonals) and genetic correlation estimates yielded from multi-trait models (described by Multi-trait model 1) using pedigree and single-step approaches.

| *Pedigree BLUP* |  |  |  |  |  |  |
| --- | --- | --- | --- | --- | --- | --- |
|  | SC | VCL | LVCL | PRS | STR | FL(m) |
| SC | **0.30** (0.06) | 0.23 (0.21) | -0.04 (0.21) | 0.18 (0.23) | 0.08 (0.22) | -0.46 (0.11) |
| VCL |  | **0.16** (0.04) | 0.84 (0.08) | 0.99 (0.04) | 0.63 (0.17) | -0.13 (0.16) |
| LVCL |  |  | **0.11** (0.03) | 0.79 (0.11) | 0.21 (0.25) | -0.13 (0.18) |
| PRS |  |  |  | **0.13** (0.04) | 0.72 (0.15) | -0.10 (0.18) |
| STR |  |  |  |  | **0.11** (0.04) | 0.03 (0.19) |
| FL(m) |  |  |  |  |  | **0.63** (0.08) |
| *single-step GBLUP* |  |  |  |  |  |  |
|  | SC | VCL | LVCL | PRS | STR | FL(m) |
| SC | **0.26** (0.06) | 0.26 (0.20) | -0.10 (0.21) | 0.21 (0.22) | 0.11 (0.28) | -0.42 (0.12) |
| VCL |  | **0.16** (0.04) | 0.83 (0.08) | 0.99 (0.06) | 0.63 (0.18) | -0.20 (0.14) |
| LVCL |  |  | **0.11** (0.03) | 0.79 (0.11) | 0.20 (0.27) | -0.22 (0.17) |
| PRS |  |  |  | **0.13** (0.04) | 0.71 (0.23) | -0.18 (0.15) |
| STR |  |  |  |  | **0.11** (0.04) | -0.05 (0.20) |
| FL(m) |  |  |  |  |  | **0.58** (0.07) |

Corresponding standard error estimates are presented in parentheses. SC: sperm concentration, VCL: curvilinear velocity, LVCL: late curvilinear velocity, PRS: proportion of rapid spermatozoa, STR: straightness of spermatozoan trajectories, FL(m): fork length of male fish. In the case of pedigree BLUP convergence was achieved after 189 iterations of which 150 used the expectation-maximization (EM) algorithm, while ss-GBLUP 181 iterations were needed of which 150 with EM.

While standard errors of genetic correlations remain largely unchanged with single-step relative to pedigree-based estimates, the direction of these associations appears more pronounced (larger absolute values), especially for correlations between kinematic parameters and either SC or FL(m). Overall, relationships between kinematic parameters appear to be strongest for the most part, while the same traits are weakly correlated with sperm concentration with the covariance estimates overlapping zero in most cases according to the corresponding standard errors. Interestingly, fork length presents mostly negative correlations with the studied male fertility proxies. The latter pattern is profound in the case of SC that exhibits a genetic correlation of approximately -0.46 (-0.42 in single-step) with FL(m).

### Genetic parameters for female fecundity and covariance with growth and male traits

To further explore genetic covariance and possible trade-offs between fertility- and growth-related phenotypes, a multi-trait model for sex-limited traits was computed (160 REML rounds; estimates in Table 2). A narrow-sense heritability of 0.37 (SE=0.10) was estimated for spawned egg-count. The respective estimates for sperm count, female and male fork length were aligned with the analysis described above. A likely complex genetic architecture is revealed by genetic correlations among the four traits. On one hand, a correlation of 0.76 was calculated between EC and FL(f), which reveals strongly positive covariance opposed to SC and FL(m). On the other hand, the genetic correlation estimate of 0.81 between FL(m) and FL(f) implies a possible genotype-by-sex interaction since it is notably lower than one. The rest of the across-sex correlations display relatively large standard errors but can be summarized as a negative genetic covariance of SC with the two female traits and a moderately strong correlation between EC and FL(m).

**Table 2.** Heritability (diagonals) and genetic correlation estimates yielded from a multi-trait model for two pairs of sex-limited phenotypes (Multi-trait model 2).

|  | EC | FL(f) | SC | FL(m) |
| --- | --- | --- | --- | --- |
| EC | **0.37** (0.10) | 0.76 (0.12) | -0.27 (0.23) | 0.62 (0.18) |
| FL(f) |  | **0.54** (0.08) | -0.21 (0.18) | 0.81 (0.11) |
| SC |  |  | **0.29** (0.07) | -0.44 (0.14) |
| FL(m) |  |  |  | **0.57** (0.07) |

Corresponding standard error estimates are presented in parentheses. EC: egg count, FL(f): fork length of female fish, SC: sperm concentration, FL(m): fork length of male fish.

### Hierarchical model for analysis of latent fertility parameters

The fixed effect of year appears to be a major contributor in both sexes (**Figure 2**). The between-sex correlation of these estimates was -0.28 for model 1 and -0.18 for model 2, revealing a moderate inverse relationship in terms of direction and intensity of year effects on male and female fertility. Such divergence may indicate sex-specific trade-offs in reproductive investment or differing sensitivities to annual environmental fluctuations.

**Figure 2:**
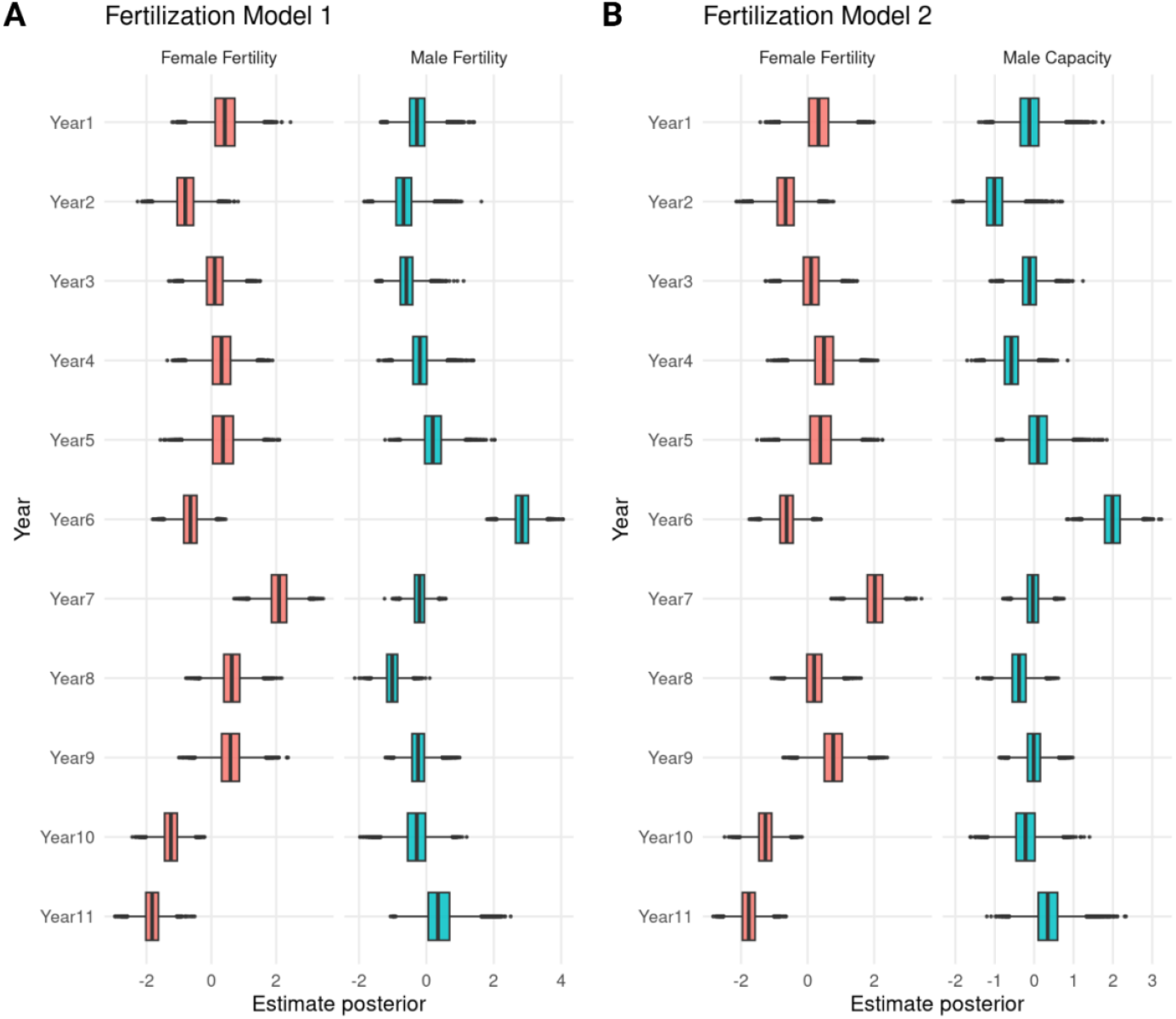
Boxplots corresponding to posterior distributions of fixed effects on latent male and female fertility traits under two hierarchical fertilization models: **A)** Fertilization Model 1 **B)** Fertilization Model 2. Figure was created with ggplot2 v3.4.4 and BioRender.com.

The variance component analysis revealed low heritability for both latent traits under Fertilization Model 1 (Table 3). Fertilization Model 2 also produced a low heritability for female fertility, which was consistent with Fertilization Model 1, but the respective estimate for male reproductive capacity was higher. In both cases, male estimates were accompanied by rather large posterior standard deviations, indicating substantial uncertainty. Weak genetic correlation estimates were obtained from both models, possibly reflecting reduced biological dependence of the traits, but reduced power due to limited data and model structure could also be a driver.

**Table 3.** Heritabilities and genetic correlations from male and female latent fertility traits.

| | Parameter | $h^2$ (sd) | $r_g$ (sd) |
| --- | --- | --- | --- |
| **Fertilization model 1** | Female fertility | 0.16 (0.05) | 0.05 (0.27) |
|  | Male fertility | 0.05 (0.04) |  |
| **Fertilization model 2** | Female fertility | 0.17 (0.04) | -0.02 (0.25) |
|  | Male capacity | 0.34 (0.20) |  |

A visualization of the estimated breeding values for latent male fertility against ssGBLUP derived proofs for observed sperm traits and fork length is provided in Figure 3. As revealed by the shape of the bivariate distributions and Pearson’s correlation coefficients (proxy genetic correlations), latent male fertility and recorded sperm phenotypes show limited association in Fertilization Model 1 (Figure 3A), with all correlations being low and/or negative. Fertilization Model 2 (Figure 3B) yields EBVs with positive correlations with key milt traits, namely sperm concentration, curvilinear velocity, proportion of rapid spermatozoa, and late curvilinear velocity. Of those, the latter seemed to be the strongest relationship and the only significant one, followed by sperm concentration. These results are expected in terms of association direction; however, the low/moderate magnitude of correlations might indicate overly simplified model assumptions or the inherently complex and potentially non-linear nature of reproductive success phenotypes, which likely reflect multiple causative biological parameters.

**Figure 3.**
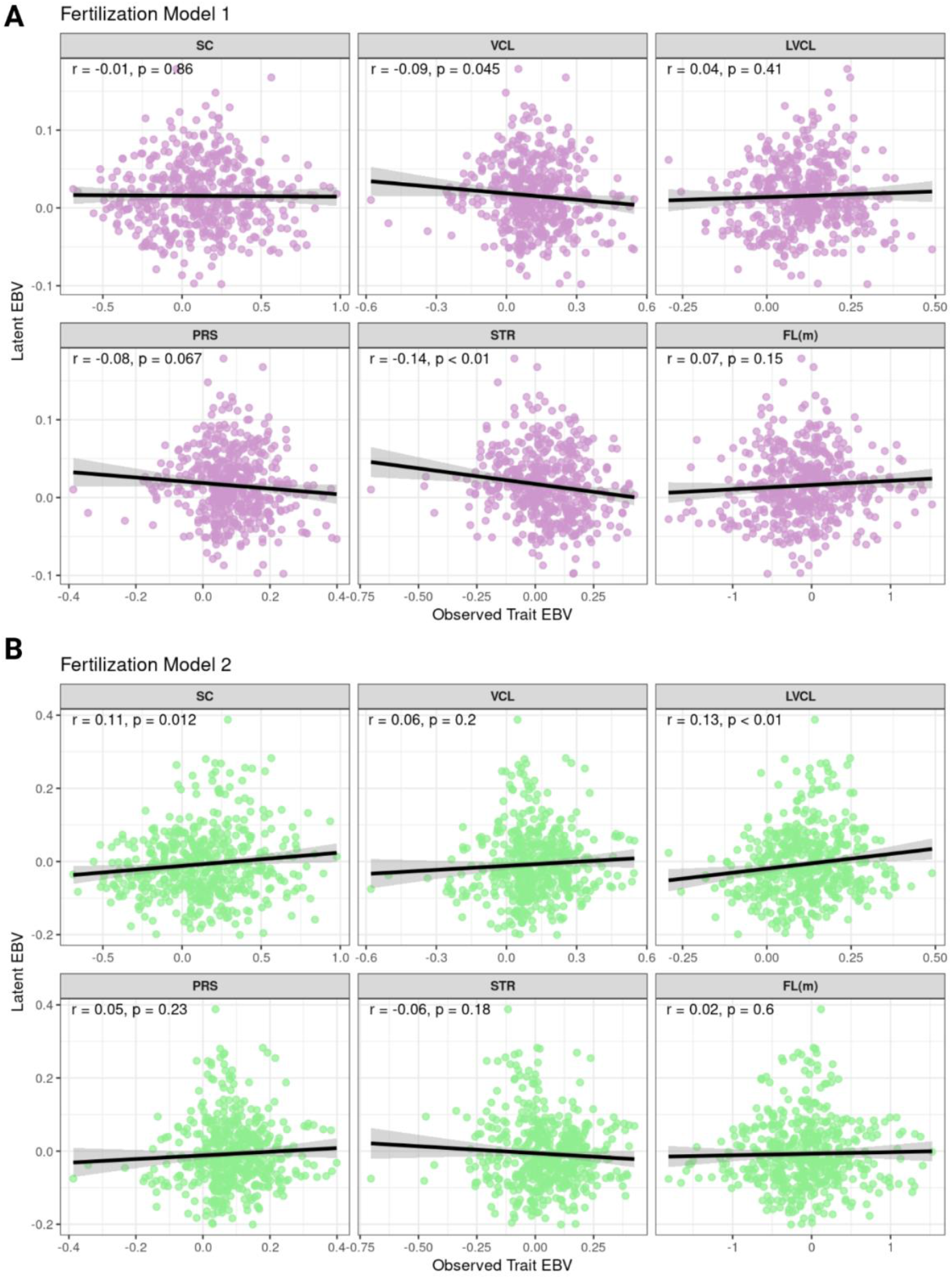
Empirical bivariate distributions of latent male fertility EBVs and observed trait EBVs under two modelling scenarios: **A)** Fertilization Model 1 and **B)** Fertilization Model 2. Corresponding regression lines, Pearson’s correlation coefficients and *p-values* are printed in all panels. SC: sperm concentration, VCL: curvilinear velocity recorded at 15 sec post-activation, LVCL: late curvilinear velocity measured at 30 sec, PRS: proportion of rapid spermatozoa, STR: straightness of trajectory. FL(m): male fork length. Figure was created with ggplot2 v3.4.4 and BioRender.com.

## Discussion

Realized fertility data possess substantial challenges due to the cryptic character of male and female contributions, non-normality of relevant data (Hartono et al., 2025) and multiple underlying genetic and environmental factors. In salmonid genetics, there is a knowledge gap about reproductive phenotypes, with the only exception being the age at sexual maturity, a trait with a strong influence on production due to the transition of energy allocation (Ahi et al., 2025; Mobley et al., 2021). Here, we estimated genetic variance components for reproductive traits with a main focus on milt-related phenotypes of Arctic charr. Additionally, a statistical framework is suggested for the genetic analysis of fertilization success rates under certain data structures and analytical assumptions.

### Genetic parameters of observed traits

In terrestrial livestock species, there is evidence for notable within- and across-species variability regarding genetic components for sperm traits, including concentration and motility; narrow-sense heritability for such traits can range from low to high depending on taxon and breed (Burren et al., 2019; Gebreyesus et al., 2021; Hu et al., 2013; Marques et al., 2017; Sá et al., 2025). In our analysis, the obtained heritability estimates for sperm quality parameters ranged from low to moderate levels (Table 1). More specifically, heritability for curvilinear velocity and proportion of rapid spermatozoa were lower than previous estimates from analysing the 8^th^ generation only, while the corresponding estimate for sperm concentration remained in line (Kurta et al., 2022). Here, we extended the aforementioned analysis by including almost thrice as many observations and included additional traits (late curvilinear velocity as a proxy for sperm motility duration; straightness of spermatozoan trajectories and fork length as a proxy for growth rate).

An interesting structure was revealed by the estimated genetic correlations between growth, sperm kinematic parameters and concentration. While motility-related parameters and straightness consistently presented positive correlations amongst them, the genetic relationship of these traits with sperm concentration seems to be generally weaker and diverging. Moreover, female fecundity appeared to negatively correlate with sperm concentration pointing towards a potential sexually antagonistic system (Table 2). This means that selection towards reproductive optima for each sex might not be possible, since this system balances around a relatively narrow overlap for male and female fertility.

Sex-specific patterns are also exposed by genetic correlations between body size and reproductive phenotypes. In females, the underlying genetic architecture for growth seems to favourably align with the one for fecundity, but the opposite is true for male where negative, low to moderate correlations with sperm phenotypes are apparent (Debes et al., 2025). This is possibly mediated by sex-specific genetic contexts for fertility (as supported by negative genetic correlations between egg count and sperm concentration). Furthermore, a genotype-by-sex interaction for growth could additionally contribute to this phenomenon. Nevertheless, sex differences in maturation age are known in Atlantic salmon (*Salmo salar*) where divergent physiology leads to earlier maturation of male individuals (Mobley et al., 2021; Tarka et al., 2018). Sexually dimorphic growth rates in favour of larger male fish have been previously reported in the studied population along with overlapping genetic signals for age at sexual maturation (Palaiokostas et al., 2022), but the present is the first attempt to assess the genetic covariance between the sexes. Based on the above findings, we hypothesize that a rather complex system is in place, characterised by an interplay of sexual antagonism and trade-offs between growth and fertility. In females, polygenic contexts for growth and fecundity appear to align, whereas in males, growth-favouring alleles may conflict with reproductive investment. This asymmetry suggests divergent selection outcomes across sexes, with clear implications for breeding strategies.

### Modelling approach for latent fertility traits

Fertilization success assessed as the ratio of eyed to spawned eggs can be thought of as a combination of three components: male fertility, female fertility and early embryo viability. Each individual component is of course a composite trait with multiple underlying biological features and can be modelled under a quantitative genetics context considering environmental and genetic contributions. Regarding the potential genetic control of early embryo viability, a major cause of mortality is catastrophic homozygosity events for deleterious alleles of genes expressed in the very early developmental stages. Such phenomena would manifest as reliably decreasing viability rates in the population by increased levels of inbreeding (Charlesworth & Willis, 2009), but such hypothesis has not been supported in the studied breeding nucleus by either pedigree-based (Palaiokostas et al., 2021) or genomics-assisted (Palaiokostas et al., 2022) investigations, likely due to lack of high inbreeding.

In contrast to traditionally applied linear models that include terms for maternal and paternal genetic contributions to reproductive outcomes (Paul et al., 2023; Piles & Tusell, 2012), we formulated hierarchical Bayesian frameworks based on two multiplicative formulas, with corresponding animal liability models as submodels for sire and dam factors. Our first proposed model considers latent female and male fertility coefficients as probabilities whose product determines the fertilization probability/rate, while the second model replaces male fertility with the complement of a Poisson void probability that accounts for the ratio between latent sperm fertilization capacity and number of spawned eggs (Figure 1B). The latter is inspired by biophysical models describing fertilization dynamics in free spawning animals, where fertilization is limited by gamete encounter rates and requires saturation by male gametes (Styan, 1998; Vogel et al., 1982). Both models assume independence of female fertility (i.e. proportion of fertile) and sperm attraction since sperm-egg interactions in this system are expected to be primarily governed by chemotaxis (Himes et al., 2011).

Notably, the variance components for the latent factors in the two models were not entirely consistent (Table 3). While the estimates from Fertilization Model 1 suggested a low heritable component dam and an even lower for sire fertility, posterior means from Fertilization Model 2 indicated moderate heritability for male capacity. Since the ground truth of female and male contributions is naturally masked, we evaluated EBVs for latent male fertility against EBVs from ssGBLUP analysis of milt phenotypes. The results of this assessment (Figure 3) implied better performance of Fertilization Model 2, since the respective proofs showed weakly positive, yet significantly correlation (*p*<0.01) with EBVs for LVCL. This might reflect possible refinement of the system due the aforementioned modelling assumptions that either ignore or include a sperm saturation effect (fertility is conditional on egg count). The exact relationships between sperm parameters and actual sire fertilizing potential are unknown. While female fertility is defined more narrowly (gamete output is known and excluded), male fertility involves multiple biological parameters, including milt volume, sperm density, motility, and durability.

An additional sex-dependent pattern was revealed by signals from environmental contributions (Figure 2), with negative correlations observed for year effects on latent male and female factors. This association implies different environmental optima for fertility between sexes. Our observation aligns with experimental evidence in Atlantic salmon supporting sex-specific response of gametes to pre-fertilization temperatures (Graziano et al., 2023). In terms of across-sex genetic correlation, estimates appeared weak under both scenarios suggesting that the genetic basis for female fertility and male reproductive capacity is likely independent. It should be stressed that the authors acknowledge the complexity of the model and recognize that parameter (including genetic variance) identifiability challenges are introduced due to latency of variables and multiple sources of variance; the components of the system are hard to disentangle, especially when paired with limited data and overdispersion driven by on-farm noise. The relatively high uncertainty in our posterior estimates probably reflects these issues. However, the synthesis of the aforementioned results, existing knowledge and consistency of variance components across models supported the robustness of the underlying trends, despite the inherent limitations.

### Relevance and future prospects

Uncovering the genetic architectures of fertilization success rates is crucial for guiding applied breeding programs and expanding fundamental biological understanding of sexual reproduction. Information about trade-offs and sexual conflict can be of great importance in designing successful breeding programs and optimizing on-farm management. In the studied nucleus, our multiplicative interpretation of fertilization rate, together with possible genotype-by-environment (G×E) interactions, may explain the apparent lack of purifying selection, despite the strong selective pressure infertility would be expected to impose. When considering the probabilistic contributions of sires and dams, even small declines in individual fertility and unfavourable genetic correlations with breeding goal traits can have a disproportionate effect on realized fertilization success rates. Under the same framework, we hypothesize that inbreeding depression in wild populations could be described more accurately: if inbreeding depression operates as pairwise multiplicative effects, reproductive success will decline disproportionately in populations with mild but widespread reductions in individual fertility, even under low incidence of strongly infertile individuals (Figure 1A). Methodological development, model optimization, simulation studies and larger-scale datasets would be needed to test such hypotheses and delve deeper into key aspects like sex-dependent G×E interactions, gamete compatibility and intrinsic effects from early embryos. Lastly, although technologies for detailed assessment of sperm quality are widely available, the standardization of reliable female fertility indicators and the integration of omic profiling of eggs are equally essential for refining and elucidating dam-specific contributions to reproductive output.

## Conclusion

Fertility is a fundamental biological characteristic of sexually reproducing animals and disruptions can lead to far-reaching consequences in breeding programs. In this study we estimated genetic variance components for reproductive proxies and growth, revealing low to moderate heritable components accompanied by a complex interplay of trade-offs and sexually antagonistic relationships. These findings suggest that conventional selection approaches may overlook important constraints, highlighting the need for more optimized breeding strategies. Additionally, we propose a modelling framework that accounts for probabilistic contributions from male and female breeders, offering an alternative approach for analysis of observed fertilization success in production settings. While the study has limitations, the compiled evidence enables inferences about sex-specific genetic and environmental effects on reproductive outcomes.

## Acknowledgements

The authors acknowledge support from the Swedish Research Council FORMAS under the EpibreedCharr project (grant agreement 2022-00348). FP was supported by the faculty of Veterinary Medicine and Animal Science, Swedish University of Agricultural Sciences.

## Author’s contributions

FP conceived methodology, performed the data analysis, and drafted the initial manuscript. CP conceived the study and, together with MJ and PVD supervised the research and edited the manuscript. All authors contributed to writing and approved the final version.

## Appendix

**Table S1:** posterior metrics and convergence diagnostics for Fertilization Model 1.

| Variable | Mean | SD | Rhat | ESS_bulk | ESS_tail |
| --- | --- | --- | --- | --- | --- |
| h2_female | 0.158 | 0.045 | 1.00 | 2348 | 4703 |
| h2_male | 0.047 | 0.043 | 1.00 | 1200 | 1388 |
| r<sub>g</sub> | 0.050 | 0.266 | 1.00 | 9938 | 14172 |

Rhat is an indicator of convergence upon approaching 1. ESS denotes the effective sample size of the Markov chain iterations. ESS\_bulk and ESS\_tail are diagnostics for the sampling efficiency in the bulk and tail of the posterior. The former is defined as the effective sample size for rank normalized values using split chains, while the latter is defined as the minimum of the effective sample sizes for 5% and 95% quantiles.

**Table S2:** posterior metrics and convergence diagnostics for Fertilization Model 2.

| Variable | Mean | SD | Rhat | ESS_bulk | ESS_tail |
| --- | --- | --- | --- | --- | --- |
| h2_female | 0.166 | 0.044 | 1.00 | 1830 | 3310 |
| h2_capacity | 0.342 | 0.199 | 1.01 | 290 | 567 |
| r<sub>g</sub> | -0.022 | 0.252 | 1.00 | 4697 | 8609 |

Rhat is an indicator of convergence upon approaching 1. ESS denotes the effective sample size of the Markov chain iterations. ESS\_bulk and ESS\_tail are diagnostics for the sampling efficiency in the bulk and tail of the posterior. The former is defined as the effective sample size for rank normalized values using split chains, while the latter is defined as the minimum of the effective sample sizes for 5% and 95% quantiles.

